# Hierarchical Reconstruction of High-Resolution 3D Models of Human Chromosomes

**DOI:** 10.1101/415810

**Authors:** Tuan Trieu, Oluwatosin Oluwadare, Jianlin Cheng

## Abstract

Eukaryotic chromosomes are often composed of components organized into multiple scales, such as nucleosomes, chromatin fibers, topologically associated domains (TAD), chromosome compartments, and chromosome territories. Therefore, reconstructing detailed 3D models of chromosomes in high resolution is useful for advancing genome research. However, the task of constructing quality highresolution 3D models is still challenging with existing methods. Hence, we designed a hierarchical algorithm, called Hierarchical3DGenome, to reconstruct 3D chromosome models at high resolution (<=5 Kilobase (KB)). The algorithm first reconstructs high-resolution 3D models at TAD level. The TAD models are then assembled to form complete high-resolution chromosomal models. The assembly of TAD models is guided by a complete low-resolution chromosome model. The algorithm is successfully used to reconstruct 3D chromosome models at 5KB resolution for the human B-cell (GM12878). These high-resolution models satisfy Hi-C chromosomal contacts well and are consistent with models built at lower (i.e. 1MB) resolution, and with the data of fluorescent in situ hybridization experiments. The Java source code of Hierarchical3DGenome and its user manual are available here https://github.com/BDM-Lab/Hierarchical3DGenome.

## Introduction

The architecture of chromosomes and genomes is important for cellular function^1,2,3^. However, the principles governing the folding of chromosomes are still poorly understood. The traditional microscopy technique - Fluorescent in Situ Hybridization (FISH) has been used to study chromosome architecture, but is limited by its low resolution and low throughput^4,5,6,7^. Chromosome conformation capture (3C) techniques like Hi-C^1^ and TCC^2^ can capture interactions between chromosomal fragments, and quantify the number of interaction frequencies (IFs) between them at a specific resolution. The bigger the interaction frequency between two fragments, the higher the probability that they are close in the three-dimensional (3D) space.

The interaction frequencies between pairs of chromosomal fragments are often summarized as a symmetric matrix, called contact matrix (or map). Contact matrices can be used to analyze the spatial organization of chromosomes or genomes. For instance, the chromosomal contact matrices have been used to confirm or identify the hallmarks of the human genome organization such as chromosome territories, chromosomal two-compartment partitions, chromatin loops, and topologically associated domains (TAD)^1,8,9^. Contact matrices can also be used to reconstruct 3D models of chromosomes and genomes to further facilitate the study of their organization. Various methods have been proposed to reconstruct 3D models of chromosomes or genomes ^10,11,12,13,14,15,16,17,18,19,20,21,22,23,24,25,26,27,28,29^. On one hand, some of these methods utilize a function that approximates the inverse relationship between interaction frequencies (IFs) and spatial distances between fragments and then uses the distances as restraints to build 3D models via spatial optimization. These methods are called the optimization based method^10,14,17^’^24^’^25^’^26^’^29^. In the early work of Duan et al.^10^, 3D models of a yeast genome were reconstructed to fit the Euclidian distances converted from IFs. On the other hand, some methods are designed to maximize the likelihood of a 3D model by using model-based methods that assumes that contact frequencies are related to distances via a probabilistic function. These methods use for example the Markov Chain Monte Carlo sampling technique to reconstruct 3D chromosome models by satisfying as many converted Euclidian distances as possible^11,12,27,28^.

Most of the existing methods are capable of reconstructing chromosome or genome models of low resolution (e.g. 1MB or 100KB) from Hi-C datasets. They can also reconstruct the 3D models of a small region of a chromosome at a high resolution. For example, LorDG^17^ built 3D models of 10 MB long chromosomal fragments at 10KB resolution. However, the ability to construct high-resolution (<=5KB) 3D models of entire large chromosomes that are needed to study the detailed interactions between genes and regulatory elements, such as enhancers at the genome scale, are still out of reach for most, if not all, of the existing methods. More recently, Rieber, L. and Mahony, S^23^, developed an approximate multidimensional scaling (MDS) algorithm called miniMDS, capable of constructing the structure of 10KB resolution from Hi-C datasets better than most of the existing methods. This algorithm partitions a Hi-C dataset into subsets, performs high-resolution MDS separately on each subset, and then reassembles the partitions using low-resolution MDS. At the time of writing this manuscript, the miniMDS is the only known method that has attempted to build a relatively higher resolution 3D models of an entire chromosome.

High-resolution genome structure modeling has several challenges. Firstly, the structure sampling is much more intensive since the number of chromosomal fragments to be modeled is much larger in high resolution. For instance, the number of chromosomal fragments to be considered at 5KB resolution is 20 times as many as at 100KB. Secondly, as the resolution increases, the number of contacts between fragments gets smaller, leaving less contact data for restraining the positions of the fragments. Finally, the search space for high-resolution models is much larger than low-resolution models, making spatial optimization much more complicated. And due to the substantial increase of the model space, models with different topologies in high resolution may satisfy the same chromosomal data. One way to reduce the search space is to require that high-resolution models have a structural topology similar to that of low-resolution models whose structure can be more stably constructed due to the availability of more contact data between larger fragments. In this work, we introduce a hierarchical algorithm to build high-resolution 3D chromosome models at 5KB resolution by using low-resolution models at 1MB resolution to assemble high-resolution models of chromosomal domains into full high-resolution models of entire chromosomes. Our results show that the high-resolution chromosome models reconstructed by our method satisfied input chromosomal contacts well, and are consistent with the experimental FISH data.

## Methods

### Data Source

We used the Hi-C contact matrices datasets (GEO accession number: GSE63525)) of cell line GM12878 ^8^ for all analyses.

### Normalization

In this work, we used two normalizations in two modelling processes. The first one is Knight-Ruiz normalization (KR) method ^30^ for building high resolution model of individual domains (step 4 in Figure 1). The second one is iterative correction and eigenvector decomposition (ICE) normalization^31^ method for building the low resolution model of the entire chromosome (step 2 in Figure 1), where each domain is represented by a point or bead.

**Figure 1.**
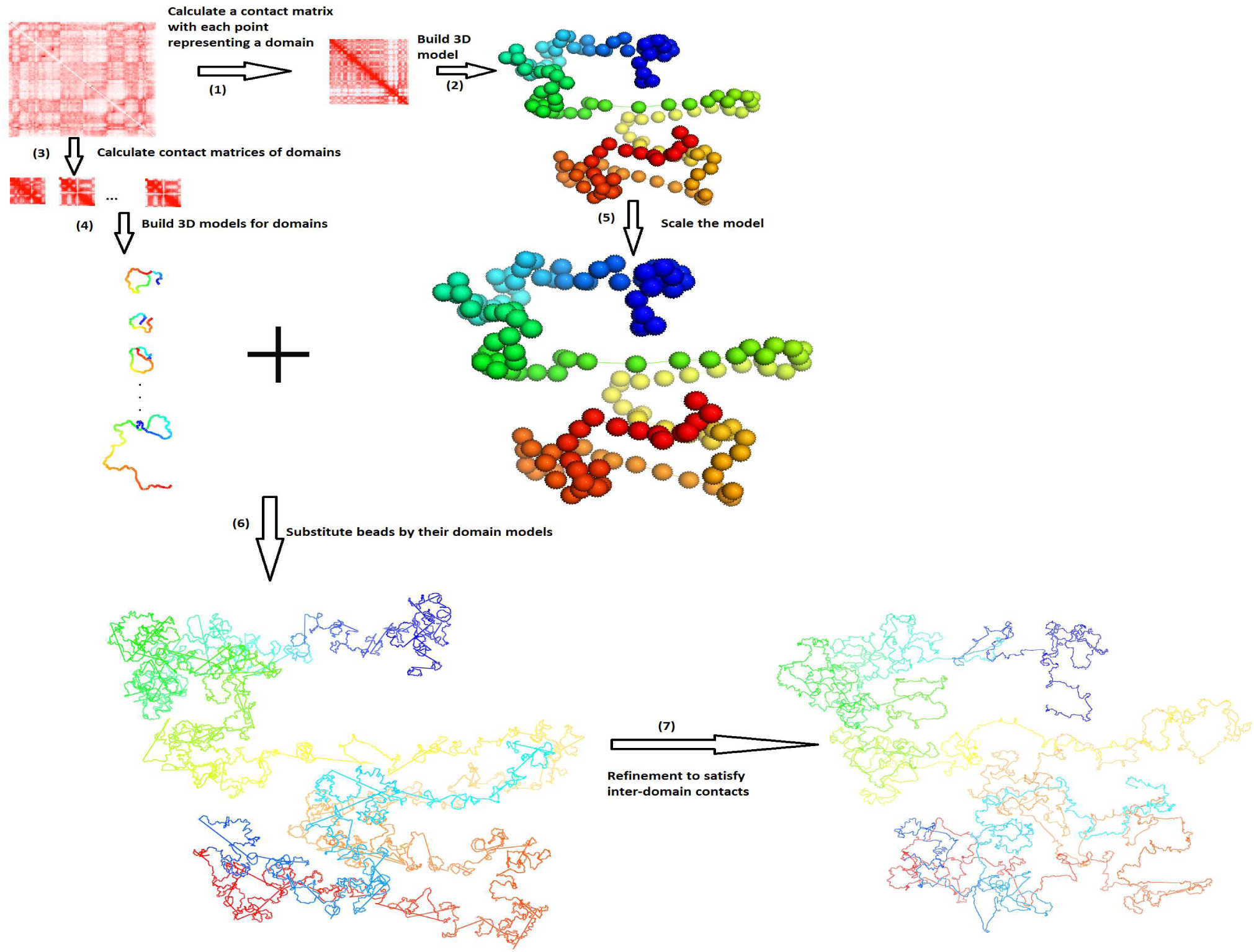
The seven steps of the hierarchical algorithm to reconstruct high-resolution models of chromosomes. The steps are: (1) Break a chromlsome into chromosomal domains according to input data and represent each domain as a point or bead, (2) Build a 3D model of the entire chromosome at low resolution, (3) Create a contact matrix for each domain, such that for n domains there are n contact matrices, (4) Build 3D model of high resolution for each domain, (5) Scale the 3D Models of the entire chromosome at low resolution to match with the models of the domains at high resolution, (6) Substitute beads in low-resolution models with their high-resolution domain models, and (7) Refine the high-resoluiton models of the entire chromosome to satisfy inter-domain contacts.

### Overview of the algorithm

A chromosome is modeled as a string of beads in 3D space, where each bead denotes the midpoint of a DNA fragment at a specific resolution (e.g. 5KB long). The position of a bead is then represented by three coordinates (*x,y,z*). Interaction frequencies between beads *i, j* are converted into spatial distances by the function 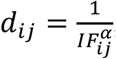, where *IF*_*ij*_ and *d*_*ij*_ are the interaction frequency and approximate distance (expected distance) between bead *i* and *j,* with *a* as a conversion factor. The goal is to place beads in 3D space so that their pairwise distances satisfy the expected distances converted from interaction frequencies as well as possible.

Our algorithm first reconstructs the 3D model of a chromosome at low resolution, which is used to guide the search for optimal models at high resolution. Each fragment (or point) in low-resolution models represents a contact domain, which is considered a structural unit of chromosome ^8^. A chromosomal domain has substantially more contacts within itself than with other domains. Therefore, the accurate models of each chromosomal domain at high-resolution can be reconstructed individually. The models of individual domains are then assembled together according to the overall topology of full chromosomal models at low resolution.

Specifically, our hierarchical algorithm, Hierarchical3DGenome, constructs high resolution chromosome 3D models in seven steps (Figure 1). The input is a contact matrix of a chromosome at a highresolution (e.g. 5KB). In Step 1, the chromosome is partitioned into contact domains (or topologically associated domains (TADs)) using the arrowhead domain algorithm ^8^. When a domain contains small domains inside, only this domain is considered because the small domains have been represented by it. Then, each separate domain is represented by a point or bead and the interaction frequencies between beads are calculated to make a low-resolution contact matrix for the entire chromosome. The matrix is normalized by the iterative correction and eigenvector decomposition (ICE) normalization ^31^ to remove technical biases ^32^, biological factors ^33^ and the different visibility of beads due to their different lengths. This new contact matrix is used to build a low-resolution model of the entire chromosome using our in-house method LorDG^17^ in Step 2. We used the default parameter settings in LorDG. The default parameter setting for LorDG algorithm allows it to search for the best conversion factor within the range [0.1, 3.0] with a step-size of 0.1 for an input contact matrix of a chromosome. The high-resolution contact matrices of individual domains are also extracted from the full high-resolution contact matrix of the chromosome in Step 3. The 3D models of each domain at the high-resolution are reconstructed individually in Step 4 (see the detailed description in the Sub Section “Construction of High-Resolution Models for each Domain”).

The topology of a correct high-resolution model of a chromosome should be similar to that of its correct low-resolution model, even though it is not guaranteed that they are in the same scale. So, in Step 5, the low-resolution model of the full chromosome constructed in Step 2 is scaled by a ratio so that it can be used to guide the assembly of the high-resolution models of individual domains into a final high-resolution model of the entire chromosome. The ratio used for this scaling is estimated from the models of individual domains and the low-resolution model of the entire chromosome (see the detailed description in the Sub Section “Estimating the Scaling Ratio between High-Resolution and Low-Resolution Models”).

After the low-resolution model is scaled to match with the scale of high resolution, in Step 6, each bead of the low-resolution model is substituted by a high-resolution model constructed in Step 4 for the corresponding domain that the bead represents. Finally, Step 7 is to further refine the location of the models of the domains to satisfy more inter-domain contacts (see the detailed description in the Sub Section “Model Refinement”).

### Construction of High-Resolution Models for Each Domain

To build high-resolution models from the contact matrix of a domain, a good conversion factor (*a)* to translate interaction frequencies into spatial distances is needed because the conversion function 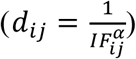 plays a crucial role in determining the quality of reconstructed models. We used the LorDG^17^ algorithm to search for the best conversion factor within the range [0.1, 3.0] with a step-size of 0.1 for an input contact matrix. Each domain could have a different conversion factor, therefore, the median of conversion factors from all the domains (e.g. *a =* 0.9) was selected as the consensus conversion factor to translate interaction frequencies into spatial distances to build high resolution models of domains with LorDG.

### Estimating the Scaling Ratio between High-Resolution and Low-Resolution Models

To estimate the ratio to scale the low-resolution model to match with the high resolution model, the distance between centers *(c*_*x*_, *c*_*y*_) of the mass of two adjacent domain models *(x, y)* is estimated by the following formula: *d*_*xy*_ *= min* _*∀ ij*_*(d*_*ij*_ *+ d*_*xi*_ *+ d*_*yj*_*·*), where *d*_*ij*_ is the distance between fragment *i* in domain *x* and fragment *j* in domain *y, d*_*xi*_ is the distance between *c*_*x*_ and *i* and *d*_*yj*_*·*is the distance between *c*_*y*_ and *j* (Figure 2). The rationale is that *d*_*xy*_ is always less than or equal to *d*_*ij*_ *+ d*_*xi*_ *+ d*_*yj*_. Therefore, *d*_*xy*_ can be well approximated by *min* _*∀ ij*_*(dij + d*_*xi*_ *+ d*_*y*_*D* given a sufficient number of fragments i and *j* are tested. It is worth noting that the adjacent domains are chosen because they have a high enrichment of inter-domain contacts between domains, hence, the distance estimated between them is more reliable.

**Figure 2.**
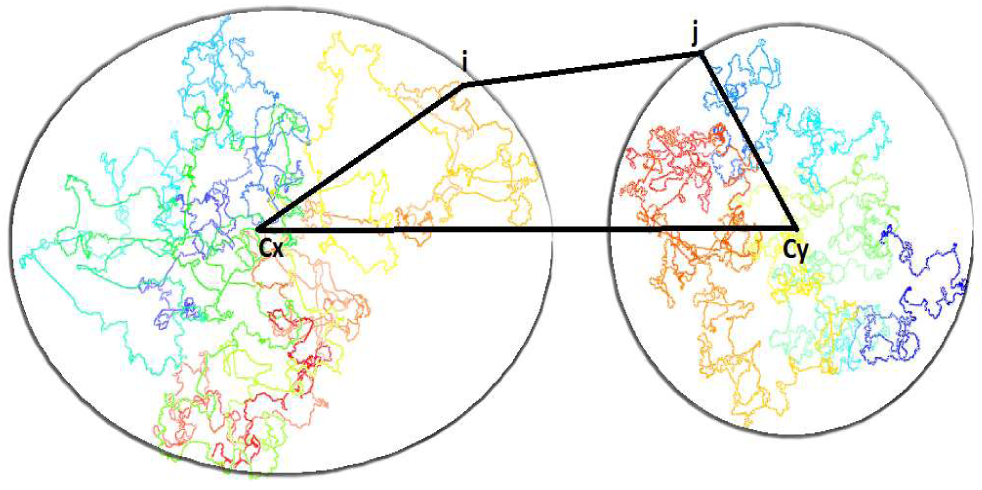
Etimation of the distance between the two centers of the two domain models

Given the 3D models of domains, the distances *d*_*xi*_ and *d*_*yj*_ can be calculated from the coordinates of the centers (i.e. the average of the coordinates of the domain model *(x,y)),* the fragment *i* in domain *x*, and the fragment *j* in domain *y*. The distance *d*can be calculated from the formula 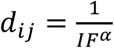 using the interaction frequency *(IF)* between fragments i and j according to the conversion factor (a) found in Step 4.

The distance between the centers of two adjacent domains, *d*_*xy*_, calculated above, are then divided by their corresponding distance in the low-resolution model to obtain a scaling ratio. In total, there will be *n* – 1 estimated distance ratios where *n* is the number of domains or beads in the low-resolution model. The final ratio used to scale the low-resolution model is the median of these estimated ratios. The centers of mass of the high-resolution domain models are placed at the locations of the corresponding beads (or points) of the low-resolution model in order to obtain an initial high-resolution model of the entire chromosome for further refinement.

### Model Refinement

In the refinement step (Step 7), we used the LorDG algorithm to adjust the coordinates of all the points of the initial high-resolution models of all the domains to satisfy high-resolution chromosomal contacts. Starting from the initial, unrefined model produced in Step 6, both intra-domain and inter-domain contacts are used in the optimization to refine it. LorDG uses all contacts to adjust the model to maximize the satisfaction of the contacts. The objective function of LorDG is non-convex and its optimization converges at local optimums. Therefore, the intra-domain contacts that have higher interaction frequency than inter-domain contacts and have already been well satisfied in the initial model are mostly preserved during the optimization. The optimization in the refinement step mostly tries to satisfy more inter-domain contacts to assemble domain models together.

## Results

We used the high-resolution Hi-C dataset^8^ with our method to build and evaluate the chromosome models at 5KB resolution. Figure 3 shows a 3D model of Chromosome 11 represented by 26,065 points at 5KB resolution, which is a 3D model of a human chromosome of the highest resolution to the best of our knowledge. We conducted five tests to evaluate the quality of these models. Firstly, we calculated the correlation between the fragment-fragment distances in the models and the expected distances calculated from contact matrices. Secondly, we checked if our 5KB-resolution models were consistent with models reconstructed directly from contact matrices at 1MB resolution. Thirdly, to figure out if domain models were well adjusted to satisfy inter-domain contacts, we extracted contact sub-matrices for every two adjacent domains consisting of both inter- and intra-domain contacts, reconstructed 3D models of the two adjacent domains from the matrices, and then compared them with the corresponding models of the two domains extracted from the full-chromosome models at 5KB resolution. Fourthly, we investigated if the high resolution chromosomal models were consistent with the FISH data^8^. The comparison shows that our models exhibited the 4 loops on four different chromosomes that were identified from the FISH data. Finally, we compared our method with an existing method for high-resolution modeling.

**Figure 3.**
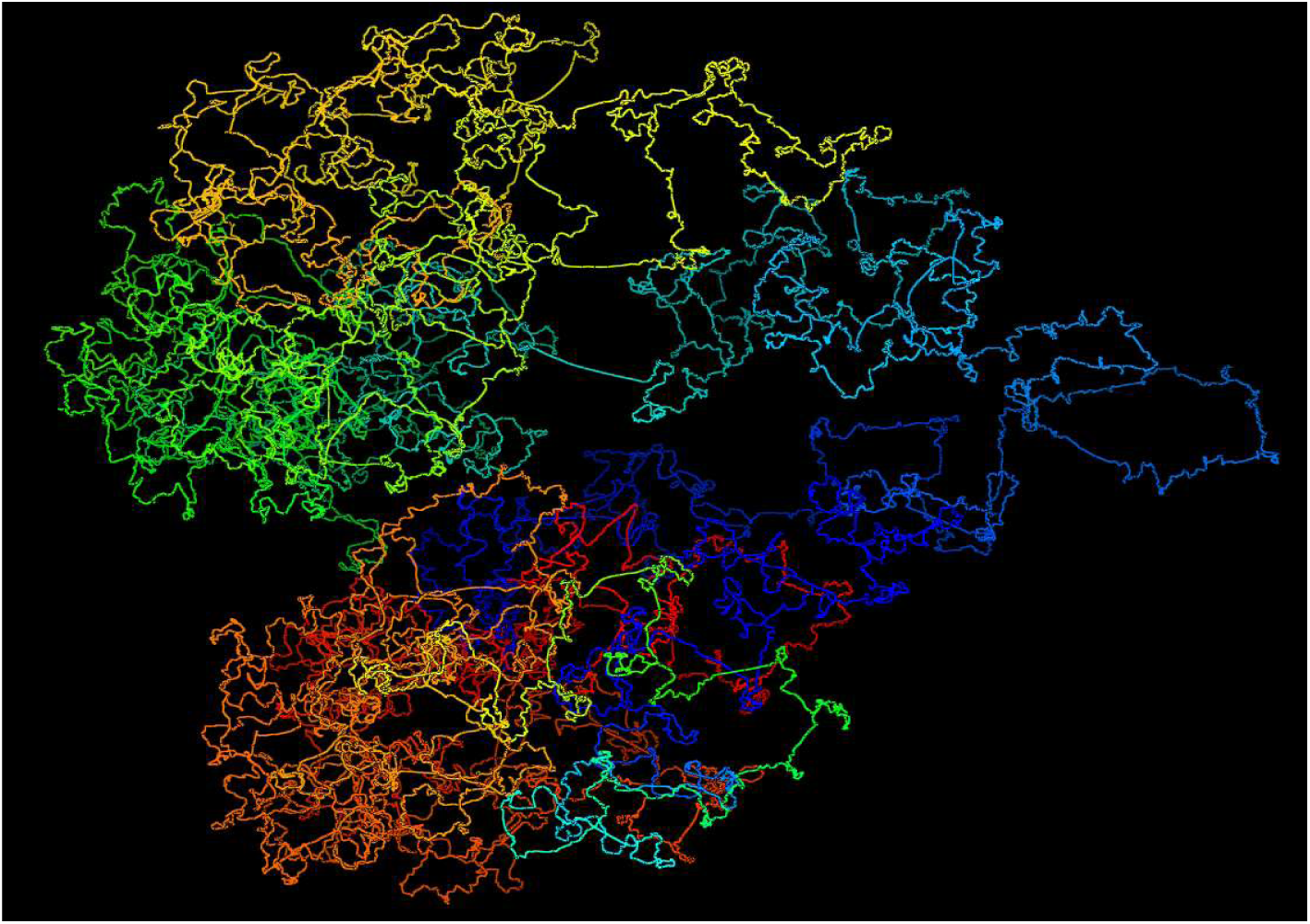
A 3D strucutre of Chromosome 11 of the cell line GM12878 at 5KB resolution

### Correlation between Reconstructed Distances and Expected Distances

We calculated the Spearman’s correlation between reconstructed fragment-fragment distances in the 3D models and their expected distances derived from the input contact matrices (Figure 4). The average and standard deviation of the correlations are 0.5357 and 0.0397, respectively. Considering the large number of expected distances to be correlated for each chromosome (e.g. 42,000,000 expected distances for Chromosome 1) and a lot of noise and inconsistency in these distances, these correlation values are good and suggest the 3D structures reconstructed models are of reasonable quality.

**Figure 4.**
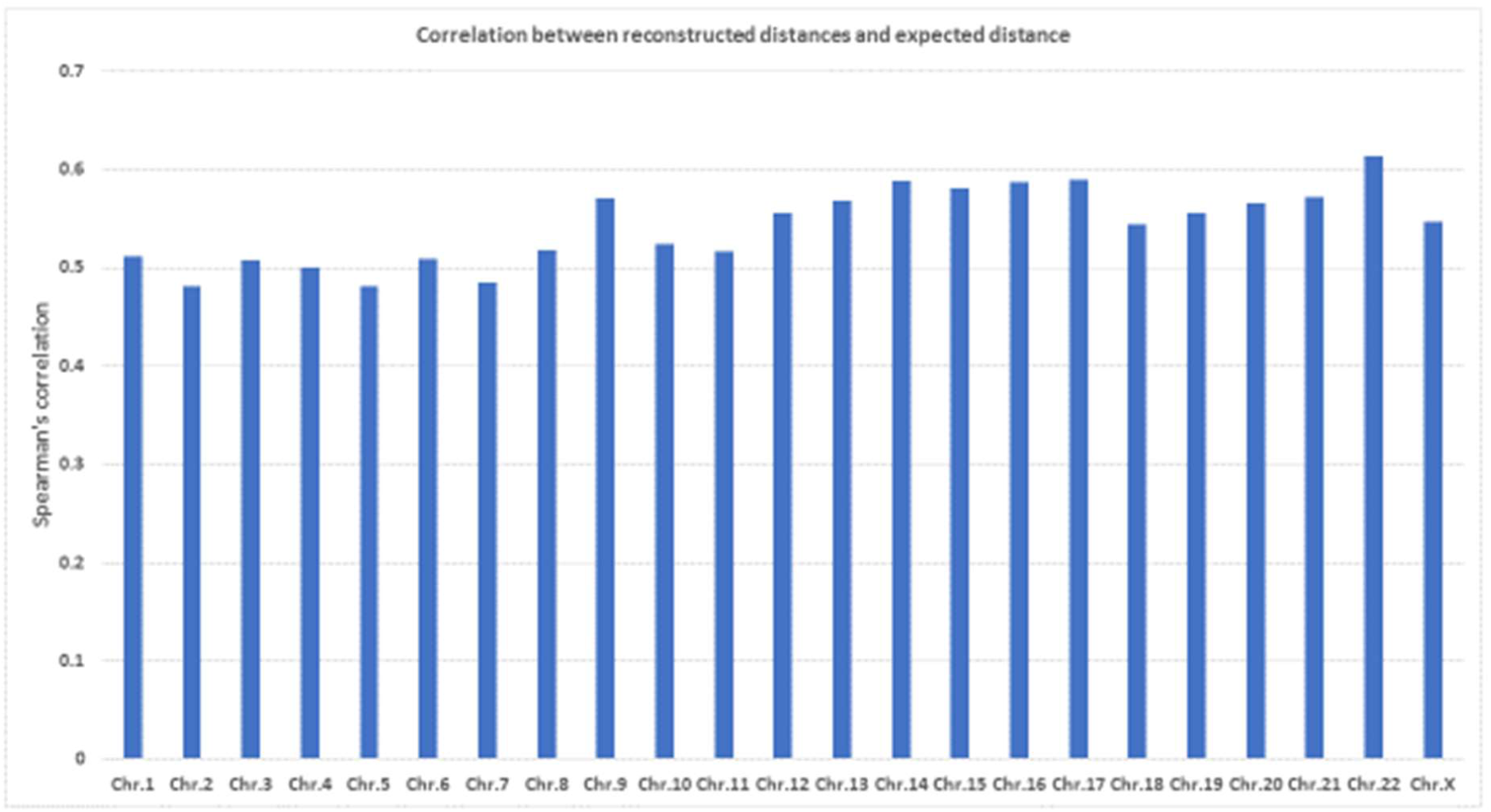
Correlation between reconstructed distances and expected distances for all 23 pairs of human chromosomes.

### Consistency between models of 5KB resolution and 1MB resolution

At 1MB resolution, because there are more restraints (reads data) between points (chromosomal fragments), the topology of models is generally more stable. We compare 5KB-resolution models with the models built directly from contact matrices of 1MB resolution to check if their topologies are consistent. To make this comparison, the 5KB resolution models were zoomed out (reduced the resolution) to 1MB resolution models. This was achieved by averaging coordinates of points of the same bin which are 1MB long as in the 1MB resolution models. We then calculated Spearman’s correlation between pairwise distances of the zoomed-out model and the 1MB resolution models (Figure 5). For all chromosomes, the correlations are ≥ 0.71, suggesting that models at 1MB and 5KB have the similar topologies.

**Figure 5.**
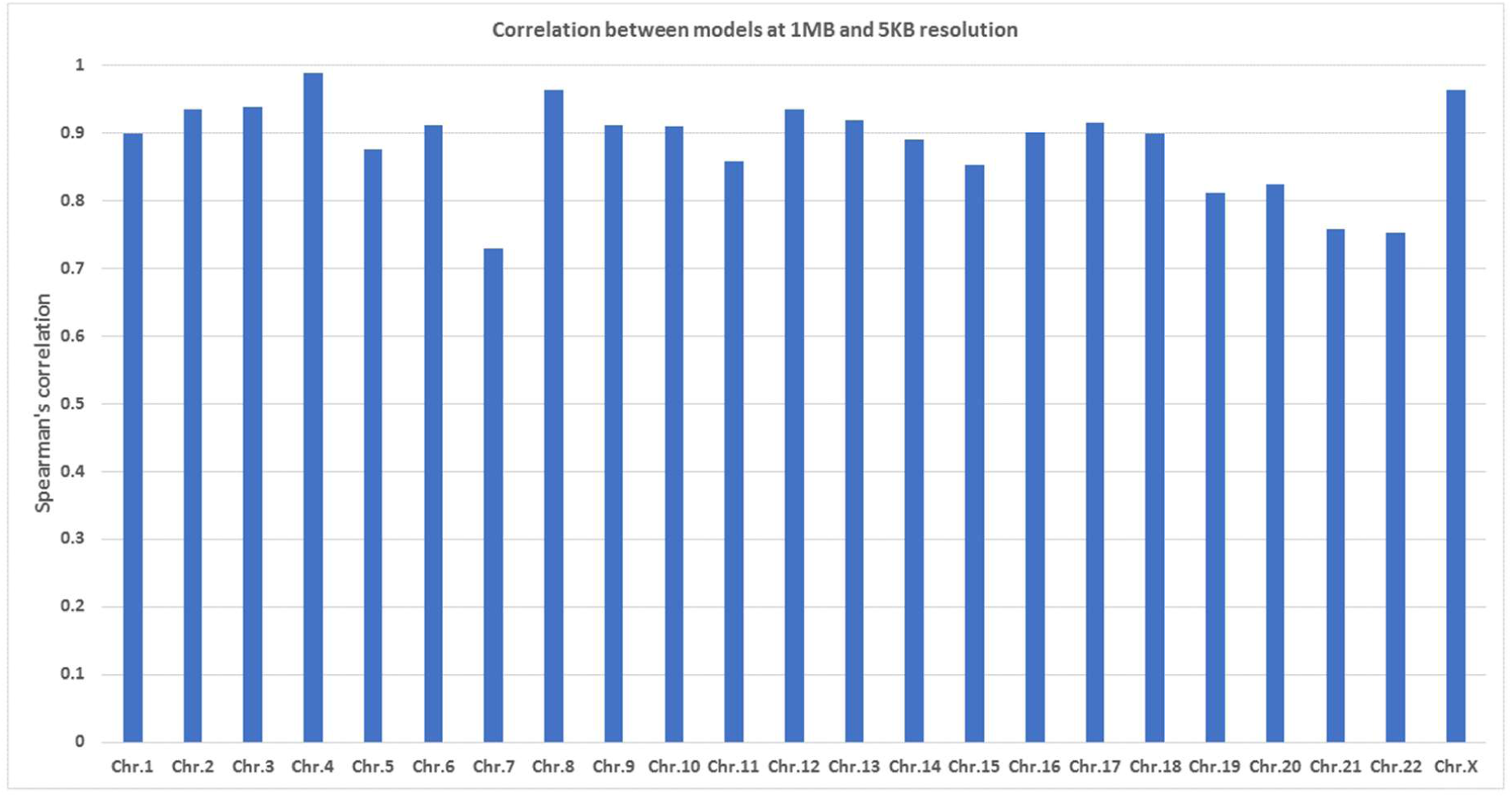
Spearman’s correlation between distances from models at 1MB and 5KB resolution. Models at 5KB resolution are consistent with models at 1MB resolution.

### Evaluation of inter-domain interactions

To check if domain models were adjusted appropriately to satisfy inter-domain contacts, we built models of every two adjacent domains from their contact sub-matrices extracted from the full contact matrix of a chromosome. We compared these models with their counterparts extracted from 5KB-resolution full-chromosome models. The boxplot in Figure. 6 shows Spearman’s correlations between the models reconstructed individually and those extracted from the 5KB resolution models for all chromosomes. The high average correlations suggest that the domains in the high-resolution full-chromosome models were generally well adjusted to satisfy inter-domain contacts.

**Figure 6.**
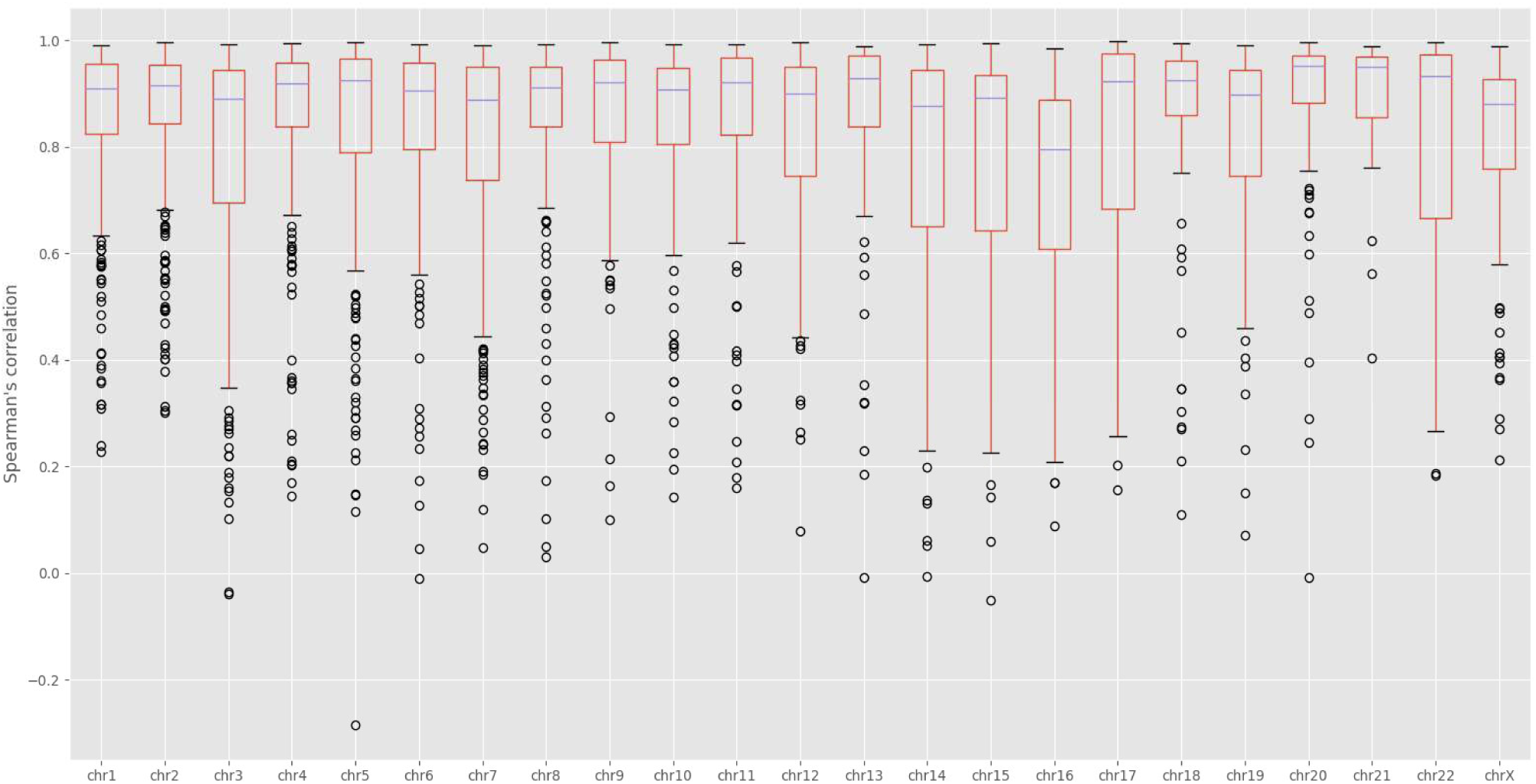
The box plot of similarity scores between models of two adjacent domains reconstructed individually and those extracted from the full chromsome model at 5KB resolution. Similarity is measured as Spearman’s correlation between distances from models. The average correlations are high. The result suggests that inter-domain contacts were well satisfied.

### Validation with FISH Data

We validated our models against the Fluorescence In Situ Hybridization (FISH) data ^8^. The FISH data identified four chromatin loops in four chromosomes of the cell line GM12878 (Chr. 11, 13, 14 and 17). For each loop, the distances between three probes (or loci), i.e., L1, L2 and L3 on the loop, were measured by FISH experiments. Although the three probes have the same genomic distance, the spatial distance of L1-L2 is much smaller than the spatial distance of L2-L3 according to the FISH experiment.

We calculated the spatial distances L1-L2 and L2-L3 in the 5KB high-resolution models and confirm that the distance of L1-L2 is indeed much smaller than the distance of L2-L3, indicating that the loops involving L1 and L2 were correctly reconstructed in the 3D models. These distances between the probes are shown in Table 1. The four loops in our models are visualized and shown in Figure. 7.

**Table 1.** The genomic positions of three probes (L1, L2, and L3) on the four loops of four chromosomes (Chr. 11, 13, 14 and 17) and the distances between these probes in the high-resolution model. Columns 2-4 list the start/end position of each probe. Columns 5 and 6 report the distances between the three probes.

| Chr. | L1 (position, MB) | L2 (position, MB) | L3 (position, MB) | L1-L2 | L2-L3 |
| --- | --- | --- | --- | --- | --- |
| 11 | 130.72-130.75 | 130.29-130.32 | 129.86-129.89 | 0.8 | 7.1 |
| 13 | 86.37-86.40 | 85.46-85.49 | 84.55-84.58 | 1.9 | 13.9 |
| 14 | 71.60-71.63 | 72.20-72.23 | 72.80-72.83 | 1.2 | 7.8 |
| 17 | 66.76-66.79 | 67.22-67.25 | 67.68-67.71 | 1.2 | 9.7 |

**Figure 7.**
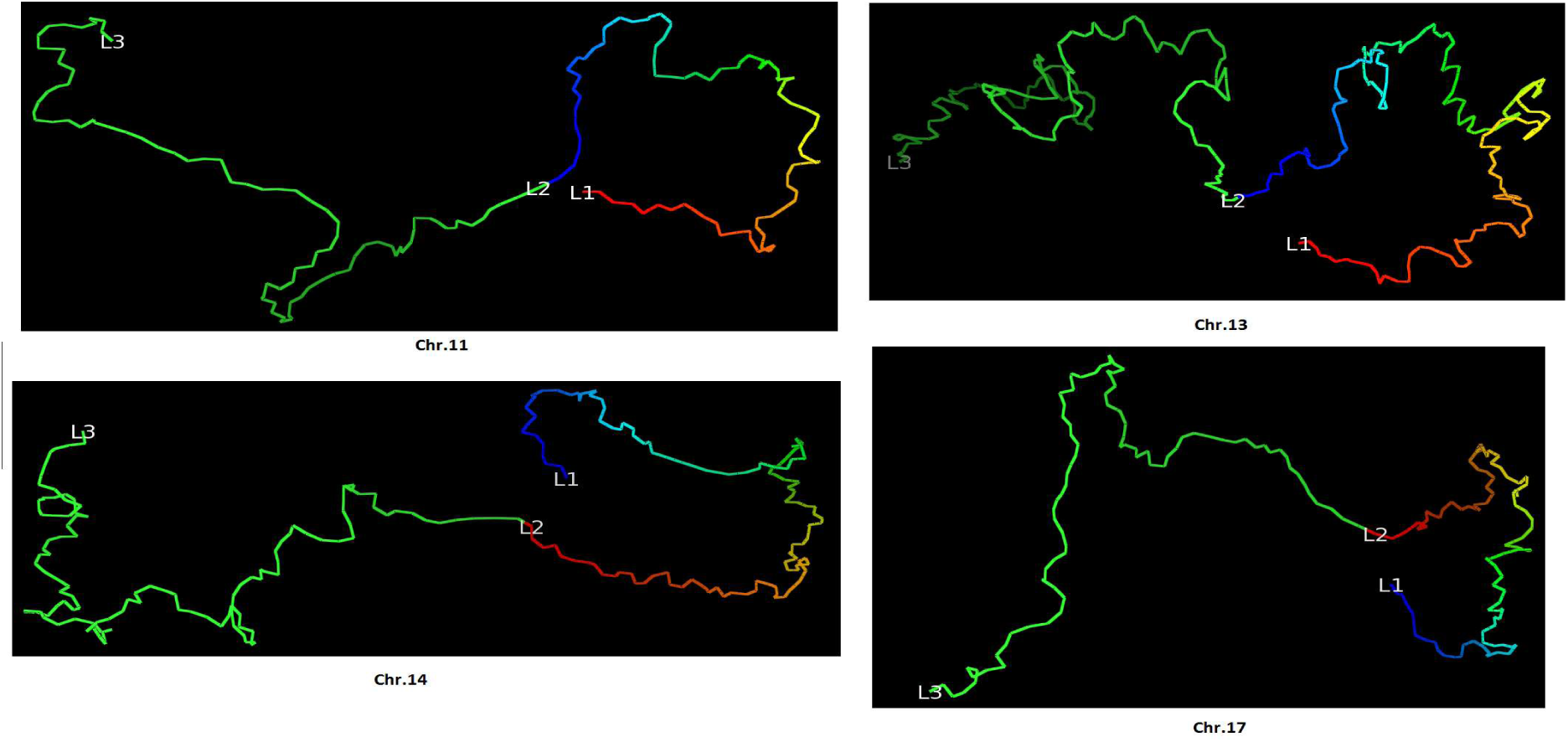
The four loops validated by FISH data for four different chromosomes (Chr. 11, 13, 14 and 17 of cell line GM12878) in the model at 5KB resolution.

### Comparison with an existing high-resolution model construction method

We compared Hierarchical3DGenome with the state of the art high resolution model construction method, miniMDS ^23^. We calculated the Spearman’s correlation between input distances and distances inferred from the output 3D structure of each chromosome at 5KB resolution produced by miniMDS. Only the non-missing points identified in the miniMDS output structure were used in this calculation. The default parameters of miniMDS were used. The result shows that the correlation is higher for Hierarchical3DGenome for every chromosome than miniMDS, indicating that Hierarchical3DGenome infers 3D structures that are more consistent with the input Hi-C data (Figure. 8). The structure from miniMDS sometimes contains folds or cluttered points with unrecognizable chromosomal features and quite some fragments are missing. In comparison, the features in the structure of Hierarchical3DGenome are well distinguished (Figure 9A, B, C versus Figure. 9D, E, F).

**Figure 8.**
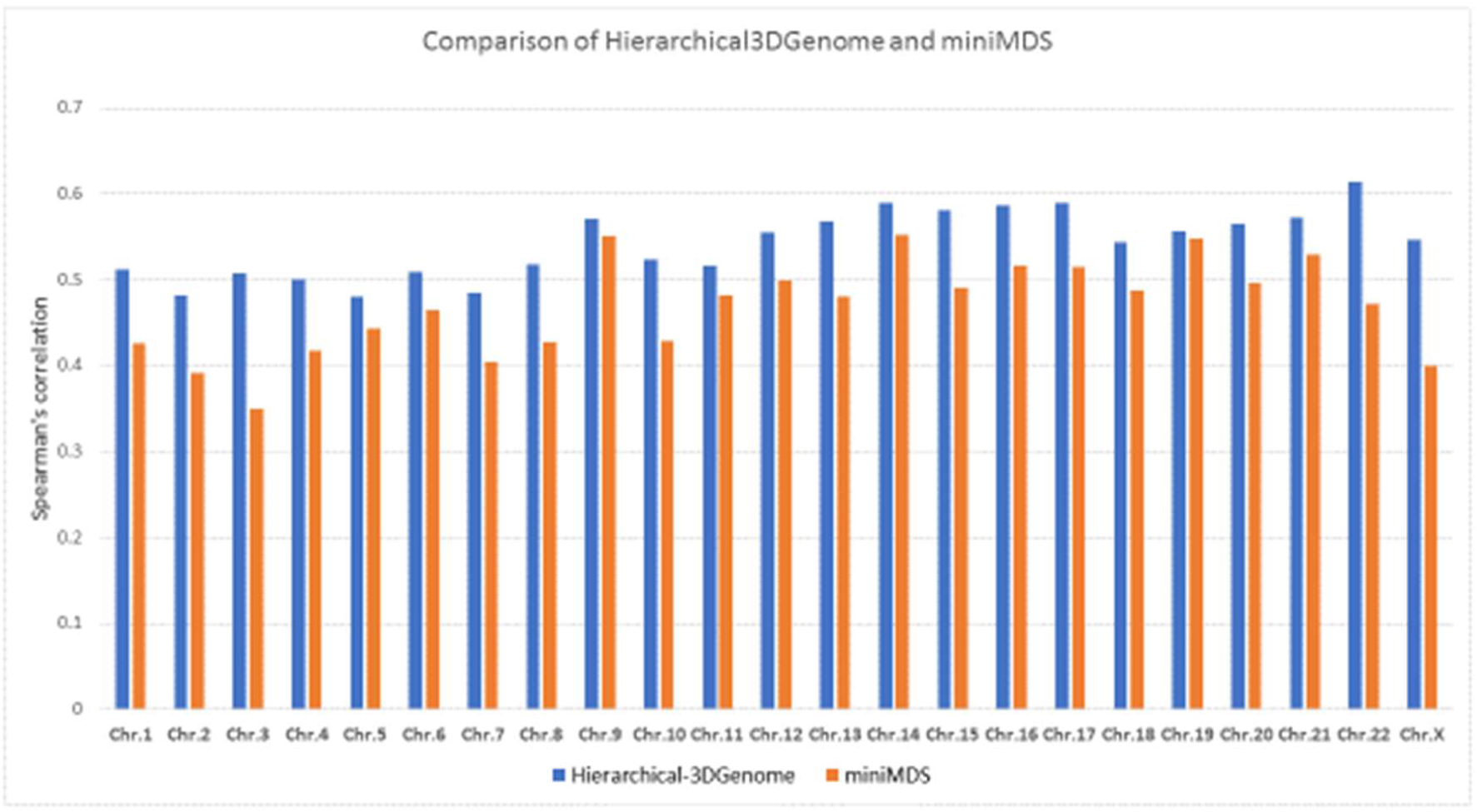
Accuray of HierarchicaBDGenome and miniMDS for each chromosome at 5KB

**Figure 9.**
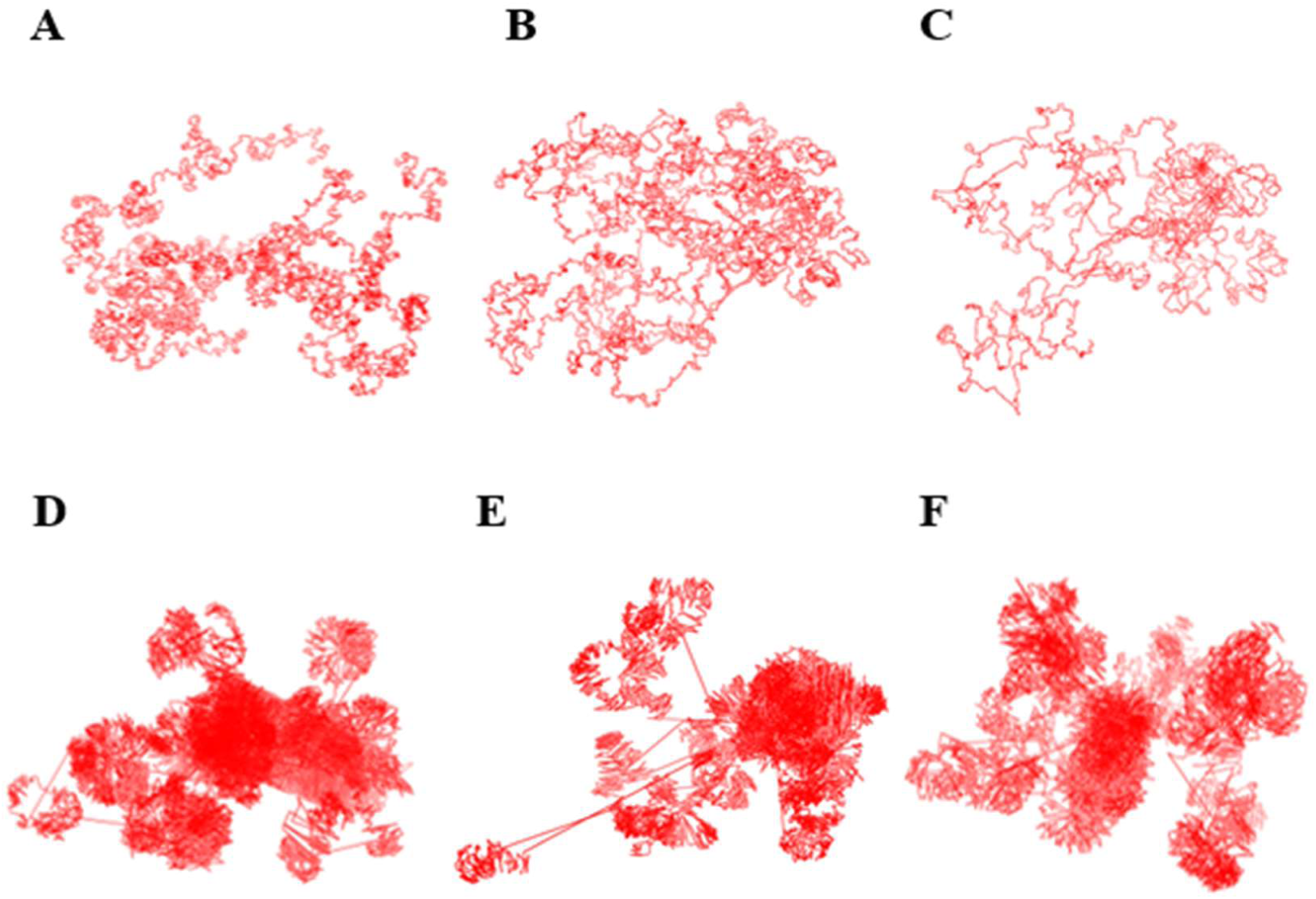
Models of Chromosomes 16,19, and 21 of 5KB resolution produced by Hierarchical3DGe-nome (A, B, C) and miniMDS (D, E, F) respectively.

### Chromosome Models at 1KB Resolution

We attempted to build chromosomal models at 1KB resolution for Chromosomes 1, 11, 13, 14 and 17 to test the capability of our method for building models of even higher resolution (Figure. 10). These models satisfied input contacts reasonably well. Spearman’s correlations between expected distances and reconstructed distances were 0.44, 0.45, 0.43, 0.48 and 0.53 respectively for these chromosomes. They were also similar to the models at 1MB resolution, indicated by a high Spearman’s correlations (> 0.89) between them. However, the detailed shapes of the four loops identified by FISH experiments on chromosome 11, 13, 14 and 17 [8] in the models of 1KB resolution did not appear as loop-like as they did in the models of 5KB resolution. One possible reason is that the input chromosomal contact data is not dense enough to build the loops of 1KB resolution, which were initially predicted with HiC data at 10KB resolution prior to being verified by FISH experiments. Another reason could be that at 1KB resolution, the increased level of noise and structural variance in the dataset reduced the prediction performance of the basic modeling tool (e.g. LorDG) used by the hierarchical modeling algorithm in this work. The problem in the former case can be solved if the Hi-C data of higher quality and resolution can be generated. In the latter case, the hierarchical modeling algorithm could be further improved by using a more robust, basic 3D modeler to build domain models. The two issues will be investigated in the future.

**Figure 10.**
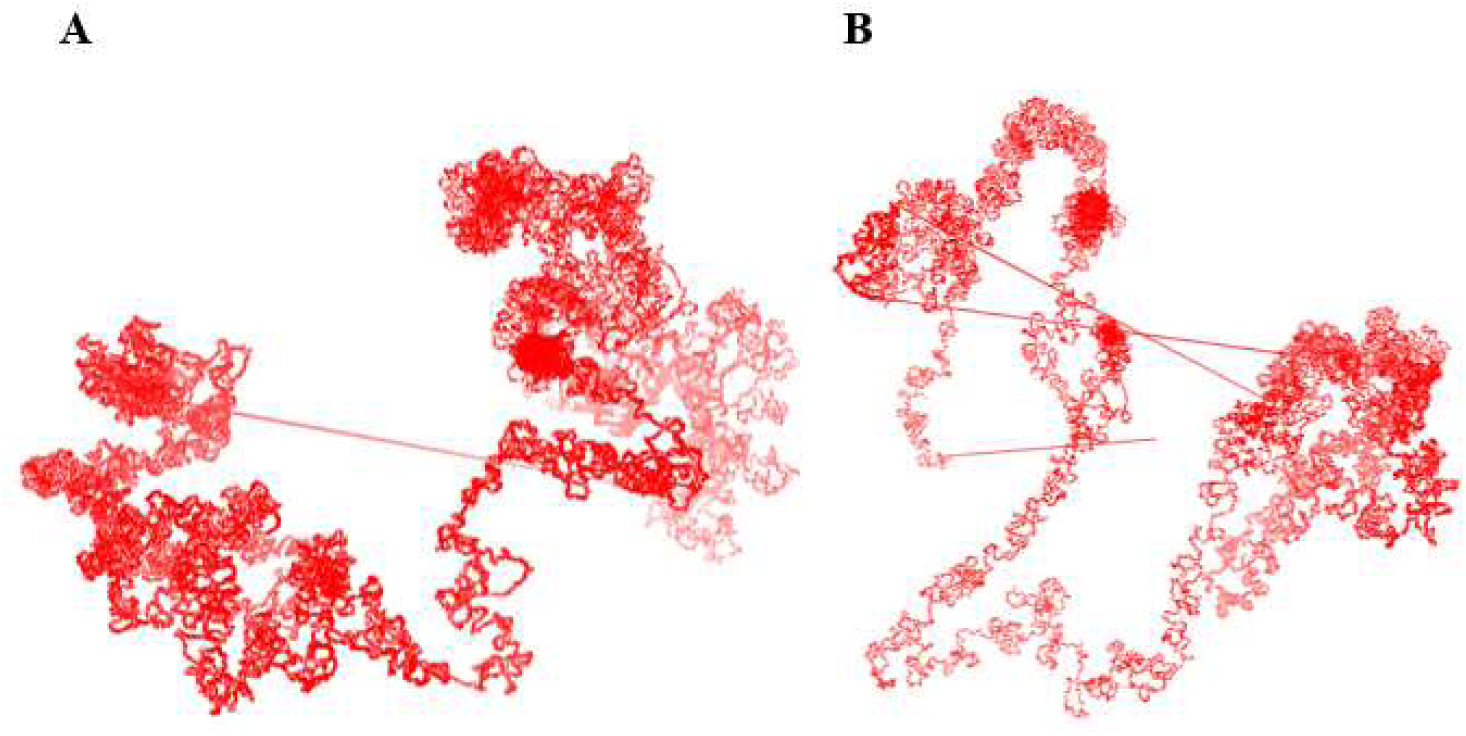
1KB resolution structures of Chromosome 1 (A) and Chromosome 11 (B) produced by Hierarchical3DGenome.

## Discussion

We introduced a hierarchical modeling algorithm to build high-resolution models of chromosomes by using low-resolution chromosomal models as a framework to position high-resolution models of topologically associated domains (TADs) constructed from Hi-C data. The algorithm was able to successfully reconstruct 3D models of full chromosomes of the human cell line GM12878 at 5KB resolution. These high-resolution models were consistent with models at 1MB resolution and FISH data. This algorithm has the potential to reconstruct 3D chromosome models at even higher resolution given HiC data of sufficient sequencing depth and quality.

## Acknowledgements

This work has been supported by National Science Foundation [grant no: DBI1149224 to JC].

## Competing interests

The authors declare no competing financial interests.

## Authors’ contributions

JC conceived the idea and supervised the project. TT designed and implemented the method. TT and OO, performed the experiments and analyzed the results. TT, OO, and JC wrote and edited the manuscript.

## Data availability

The Java source code of Hierarchical3DGenome, sample datasets, its user manual, and the parameters for running the different analysis are available here: https://github.com/BDM-Lab/Hierarchical3DGenome

## Acknowledgements

None

